# Harnessing the FBXW7 somatic mutant R465C for targeted protein degradation

**DOI:** 10.1101/2024.12.03.626601

**Authors:** Ananya A. Basu, Chenlu Zhang, Milad Rouhimoghadam, Anil Vasudevan, Justin M. Reitsma, Xiaoyu Zhang

## Abstract

Targeted protein degradation (TPD) is a pharmacological strategy that eliminates specific proteins from cells by harnessing cellular proteolytic degradation machinery. In proteasome-dependent TPD, expanding the repertoire of E3 ligases compatible with this approach could enhance the applicability of this strategy across various biological contexts. In this study, we discovered that a somatic mutant of FBXW7, R465C, can be exploited by heterobifunctional compounds for targeted protein degradation. This work demonstrates the potential of utilizing mutant E3 ligases that occur exclusively in diseased cells for TPD applications.

---

Targeted protein degradation (TPD) is an emerging strategy that employs small molecules or biologics to direct proteins to proteolytic degradation machinery, such as the proteasome and lysosome, facilitating the removal of target proteins^1, 2^. In ubiquitin–proteasome system (UPS)-dependent TPD, heterobifunctional compounds, known as PROTACs (proteolysis-targeting chimeras)^3^, or monofunctional molecular glues^4^, induce proximity between the target protein and an E3 ligase. This allows the E3 ligase to ubiquitinate the target protein, directing it for subsequent degradation by the proteasome. The E3 ligase family comprises over 600 members in humans, yet most TPD-susceptible proteins identified to date are degraded through two: CRBN and VHL^2, 5^. While CRBN and VHL are widely utilized in TPD, expanding the repertoire of TPD-compatible E3 ligases has the potential to broaden the applicability of this strategy across diverse targets and diseases.

Drugs that selectively target mutant forms of proteins can differentiate between proteins in healthy and diseased cells, thereby enhancing selectivity in disease cells. Analyses of genome-wide mutational spectra in human cancers reveal that approximately 17% of all missense mutations result in cysteine substitutions^6, 7^. These acquired cysteines present an opportunity for cancer-specific targeting. An example is the KRAS-G12C mutation, against which Sotorasib and Adagrasib have recently received FDA approval for patients with *KRAS*^G12C^ mutant non-small cell lung cancer (NSCLC)^8, 9^. In TPD, hijacking acquired cysteines in E3 ligases for target degradation could potentially induce target degradation in disease cells harboring these mutations, while leaving normal cells unaffected. In this work, we discovered that the FBXW7 somatic mutant R465C can be hijacked by heterobifunctional compounds for targeted protein degradation. This finding demonstrates a novel approach of harnessing mutant E3 ligases for TPD, offering potential disease-specific therapeutic advantages.

To investigate acquired cysteines in E3 ligases—some of which may exist in a reactive state amenable to ligand discovery—we examined a set of 680 human E3 ligases and analyzed 24,834 acquired cysteines occurring in 11,353 proteins cataloged in The Cancer Cell Line Encyclopedia (CCLE)^10^ (**Table S1**). This analysis revealed 1,102 acquired cysteines present on 424 E3 ligases (**Figure 1a** and **Table S1**). By further filtering for hotspot mutations from The Cancer Genome Atlas (TCGA), we identified 39 acquired cysteines in 35 E3 ligases (**Figure 1a** and **Table S1**). Ranking these acquired cysteines by mutation frequency showed that the top two, R465C and R505C, are found in the same E3 ligase, FBXW7 (**Figure 1b**). FBXW7, a member of the F-box protein family, functions as a component of the Cullin-RING ubiquitin ligase 1 (CRL1) complex^11^. FBXW7 is crucial in regulating cell survival, proliferation, tumor invasion, DNA repair, and genomic stability, establishing its importance in oncogenesis^12^. The R465 and R505 hotspot mutations are located in the WD40 domain and were initially thought to cause loss-of-function effects^13^. Since R465C is a more prevalent acquired cysteine mutation according to CCLE hotspot data and is associated with a worse prognosis in colorectal cancer patients compared to other frequent FBXW7 mutations^14^, we focused on this mutant for degrader discovery. In previous studies, we applied a “degradation-first” approach to screen a focused FKBP12-directed bifunctional compound library, aiming to identify TPD-competent E3 ligases^15-17^. Here, we first knocked out *FBXW7* in HEK293T cells (**Figure S1a**), then stably expressed FLAG-tagged nucleus-localized FKBP12 (FLAG-FKBP12_NLS) along with either wild-type (WT) or R465C mutant HA-FBXW7. As the major FBXW7 isoform is nuclear^18^, we utilized nucleus-localized FKBP12 for the screen—a strategy previously used to identify nuclear-localized TPD-competent E3 ligases^17^. We treated the cells with 22 FKBP12-targeting bifunctional compounds previously synthesized^15, 17^, and measured FKBP12 abundance in cells expressing either WT or R465C FBXW7 by Western blot. The results indicate that one compound, 10-SLF, selectively reduced FKBP12 levels in cells expressing FBXW7-R465C, but not in those expressing WT (**Figure 1c** and **Figure S1b**).

**Figure 1.**
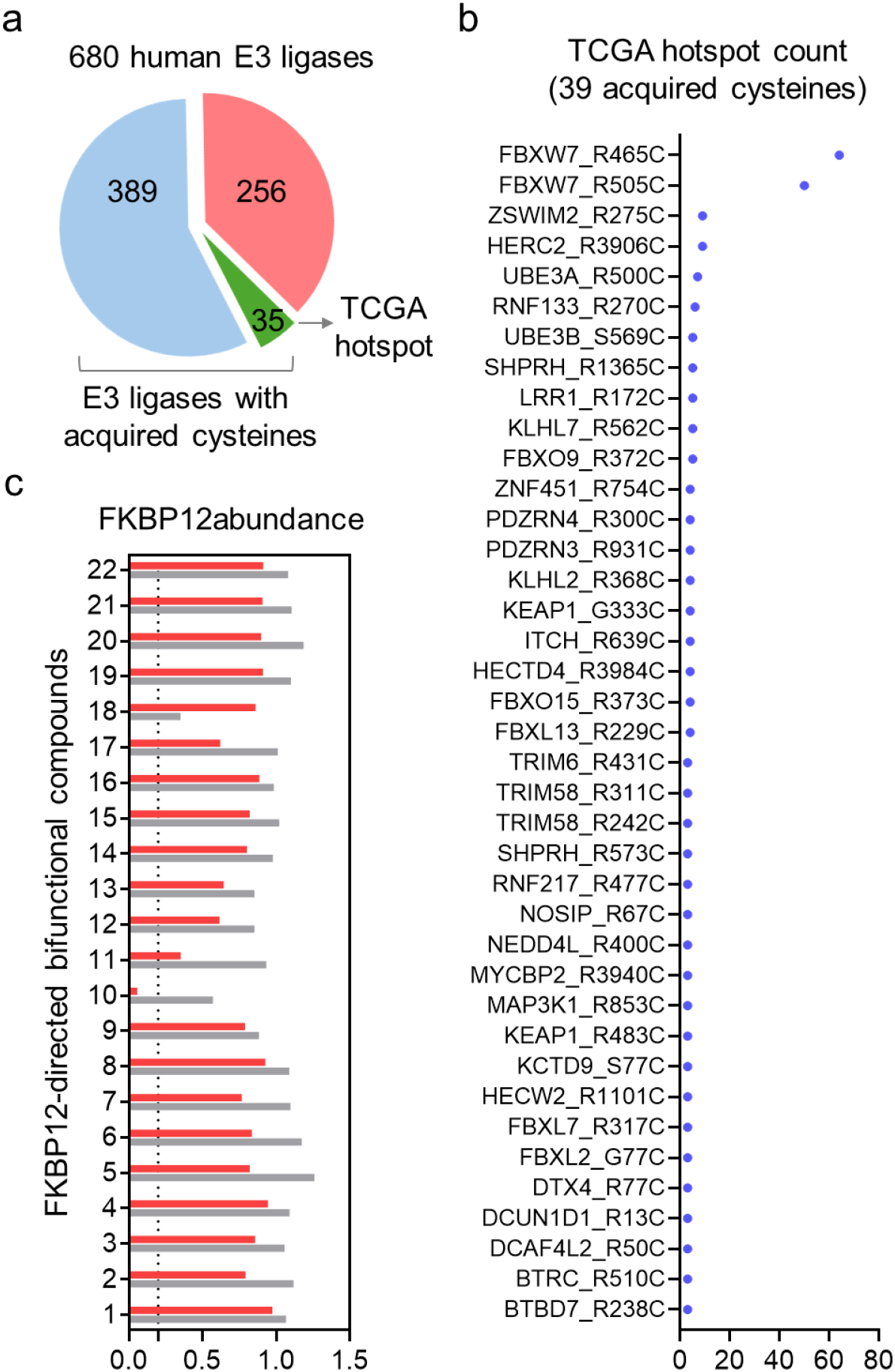
Identification of heterobifunctional compounds that degrade FKBP12 in an FBXW7-R465C-dependent manner. **a**. Analysis of acquired cysteines within 680 human E3 ligases. **b**. The TCGA hotspot count of the 39 acquired cysteines occurring in 35 E3 ligases. **c**. Screening of an FKBP12-directed heterobifunctional compound library to identify compounds that degrade FKBP12 in an FBXW7-R465C-dependent manner.

10-SLF is a bifunctional compound containing an α-chloroacetamide moiety (**Figure 2a**) that may covalently interact with FBXW7-R465C. We next tested different concentrations of 10-SLF in HEK293T cells overexpressing FLAG-FKBP12_NLS and either HA-FBXW7 WT or R465C, observing a reduction in FKBP12_NLS levels starting at 0.25 µM only in cells expressing FBXW7-R465C, but not in those expressing WT FBXW7 (**Figure 2b**). The 10-SLF-induced FKBP12_NLS reduction was blocked by MG132 (a proteasome inhibitor), SLF (a FKBP12 ligand), and MLN4924 (a neddylation inhibitor) (**Figure 2c**), suggesting that FKBP12 degradation involves both the proteasome and Cullin-RING ligase pathways, consistent with FBXW7’s role in the Cullin-RING E3 ligase family. A time-course study of 10-SLF-induced FKBP12_NLS degradation revealed maximal degradation after 8 hours (**Figure 2d**).

**Figure 2.**
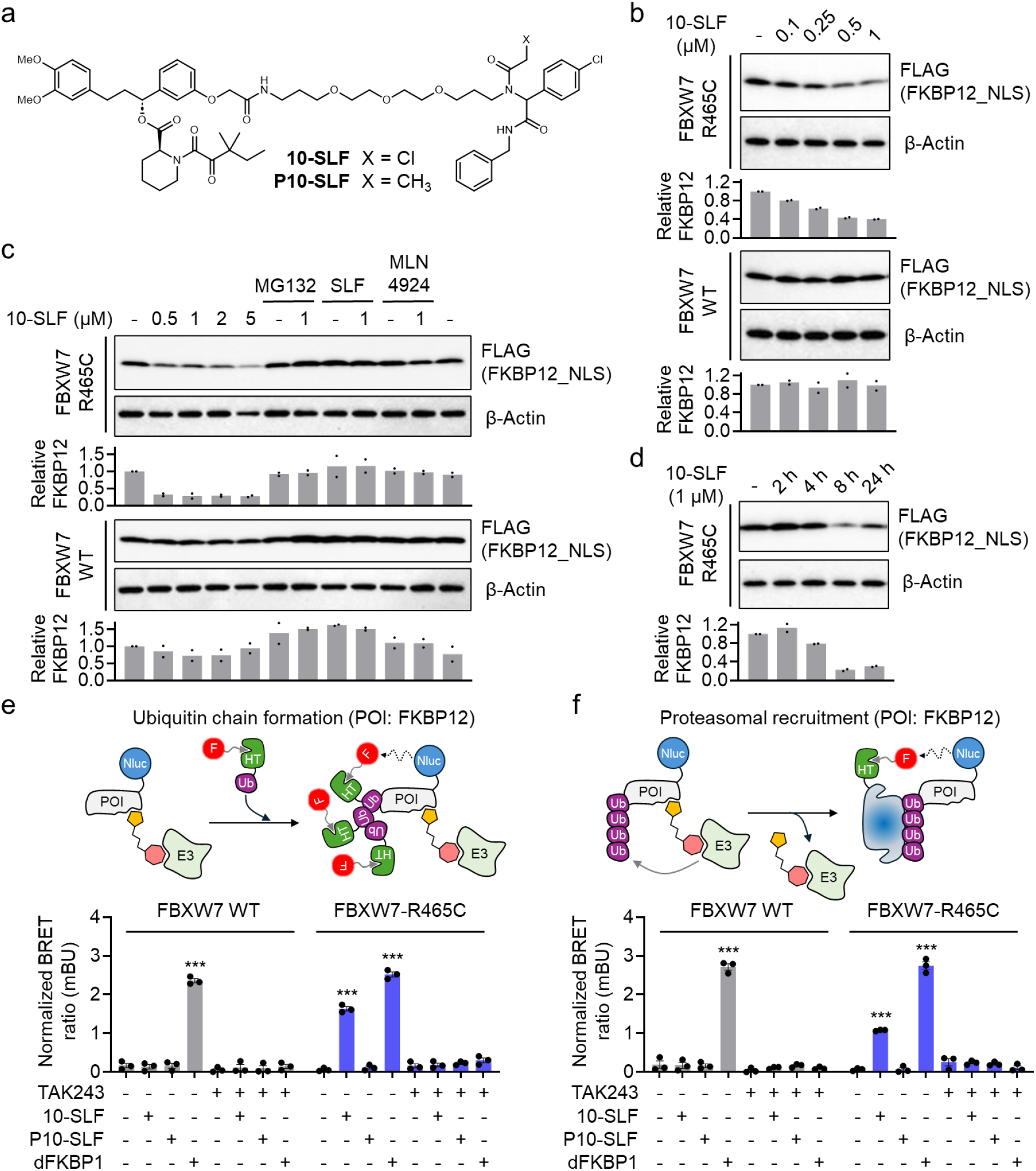
FBXW7-R465C supports 10-SLF-induced ubiquitination and degradation of FKBP12. **a**. Structures of 10-SLF and P10-SLF. **b**. Dose-dependent degradation of FKBP12_NLS by 10-SLF. HEK293T cells expressing HA-FBXW7 WT or R465C and FLAG-FKBP12_NLS were treated with 0.1–1 µM 10-SLF for 8 hours. The bar graph represents quantification of the FKBP12/*β*-Actin protein content. Data are presented as the mean values (n = 2 independent replicates). **c**. 10-SLF-induced FKBP12_NLS degradation was blocked by MG132, SLF, and MLN4924. The bar graph represents quantification of the FKBP12/*β*-Actin protein content. Data are presented as the mean values (n = 2 independent replicates). **d**. A time-course study of 10-SLF-induced FKBP12_NLS degradation. The bar graph represents quantification of the FKBP12/*β*-Actin protein content. Data are presented as the mean values (n = 2 independent replicates). **e**. NanoBRET assay measuring the ubiquitination of FKBP12 induced by 10-SLF. Data are presented as the mean values ± s.e.m. (n = 3 independent replicates). The statistical significance was evaluated through unpaired two-tailed Student’s *t*-tests, comparing cells treated with 10-SLF or dFKBP1 to DMSO. \*\*\**P* < 0.001. **f**. NanoBRET assay measuring the engagement of FKBP12 with the proteasomal submit PSMD3 induced by 10-SLF. Data are presented as the mean values ± s.e.m. (n = 3 independent replicates). The statistical significance was evaluated through unpaired two-tailed Student’s *t*-tests, comparing cells treated with 10-SLF or dFKBP1 to DMSO. \*\*\**P* < 0.001.

Next, we used two nano bioluminescence resonance energy transfer (NanoBRET) assays^19^ to assess 10-SLF-induced ubiquitination of FKBP12 and its subsequent proteasomal engagement. In the first assay, HEK293T cells stably expressing HA-FBXW7 WT or R465C were transiently transfected with nanoluciferase (Nluc)-fused FKBP12 and HaloTag-fused ubiquitin (Ub) (**Figure 2e**). Following treatment with 10-SLF and a fluorophore-conjugated chloroalkane ligand that covalently engages HaloTag-Ub, ubiquitinated FKBP12 produces a NanoBRET signal between Nluc-FKBP12 and HaloTag-Ub. In cells expressing FBXW7 WT, only the positive control PROTAC, dFKBP1^20^ (**Figure S2**), induced FKBP12 ubiquitination, whereas 10-SLF did not trigger the ubiquitination of FKBP12 (**Figure 2e**). In contrast, in cells expressing FBXW7-R465C, both dFKBP1 and 10-SLF induced significant FKBP12 ubiquitination compared to DMSO control (**Figure 2e**). We further synthesized a non-covalent analog of 10-SLF, P10-SLF, by replacing the α-chloroacetamide group with a propanamide (**Figure 2a**), which did not induce FKBP12 ubiquitination in cells expressing FBXW7-R465C (**Figure 2e**). The FKBP12 ubiquitination induced by 10-SLF or dFKBP1 was fully blocked by the E1 enzyme inhibitor TAK243^21^, indicating its dependence on the ubiquitin–proteasome system (UPS) (**Figure 2e**).

In the second assay, we assessed FKBP12 engagement with the proteasome^22^, indicating its degradation mediated by this machinery. HEK293T cells stably expressing HA-FBXW7 WT or R465C were transiently transfected with Nluc-FKBP12 and HaloTag-PSMD3, a proteasome 26S subunit (**Figure 2f**), followed by treatment with 10-SLF and a fluorophore-conjugated chloroalkane ligand. The positive control dFKBP1 induced FKBP12 engagement with the proteasome in both FBXW7 WT- and R465C-expressing cells (**Figure 2f**). Consistent with the findings in the ubiquitination assay, 10-SLF promoted FKBP12 engagement with the proteasome only in FBXW7-R465C-expressing cells, with no effect in FBXW7 WT cells (**Figure 2f**). The non-covalent control P10-SLF did not induce FKBP12 engagement with the proteasome in either cell model (**Figure 2f**), and the E1 enzyme inhibitor TAK243 blocked this engagement induced by both dFKBP1 and 10-SLF (**Figure 2f**). Collectively, these data suggest that, in the presence of FBXW7-R465C, 10-SLF induces FKBP12 ubiquitination and subsequent proteasomal degradation, with this activity requiring the compound’s covalent nature.

Next, we investigated whether endogenously expressed FBXW7-R465C supports 10-SLF-induced FKBP12 degradation. We selected AsPC-1^10^, a human pancreatic tumor cell line that expresses *FBXW7*^R465C^. We knocked out *FBXW7*^R465C^ in AsPC-1 cells (**Figure 3a**), and subsequently stably overexpressed FLAG-FKBP12_NLS to assess target degradation by 10-SLF. Our results showed that 10-SLF induced FKBP12 degradation in parental AsPC-1 cells but not in *FBXW7*^R465C^ knockout (KO) cells (**Figure 3b**). The non-covalent variant, P10-SLF, did not induce FKBP12 degradation in either parental or KO cells (**Figure 3c**).

**Figure 3.**
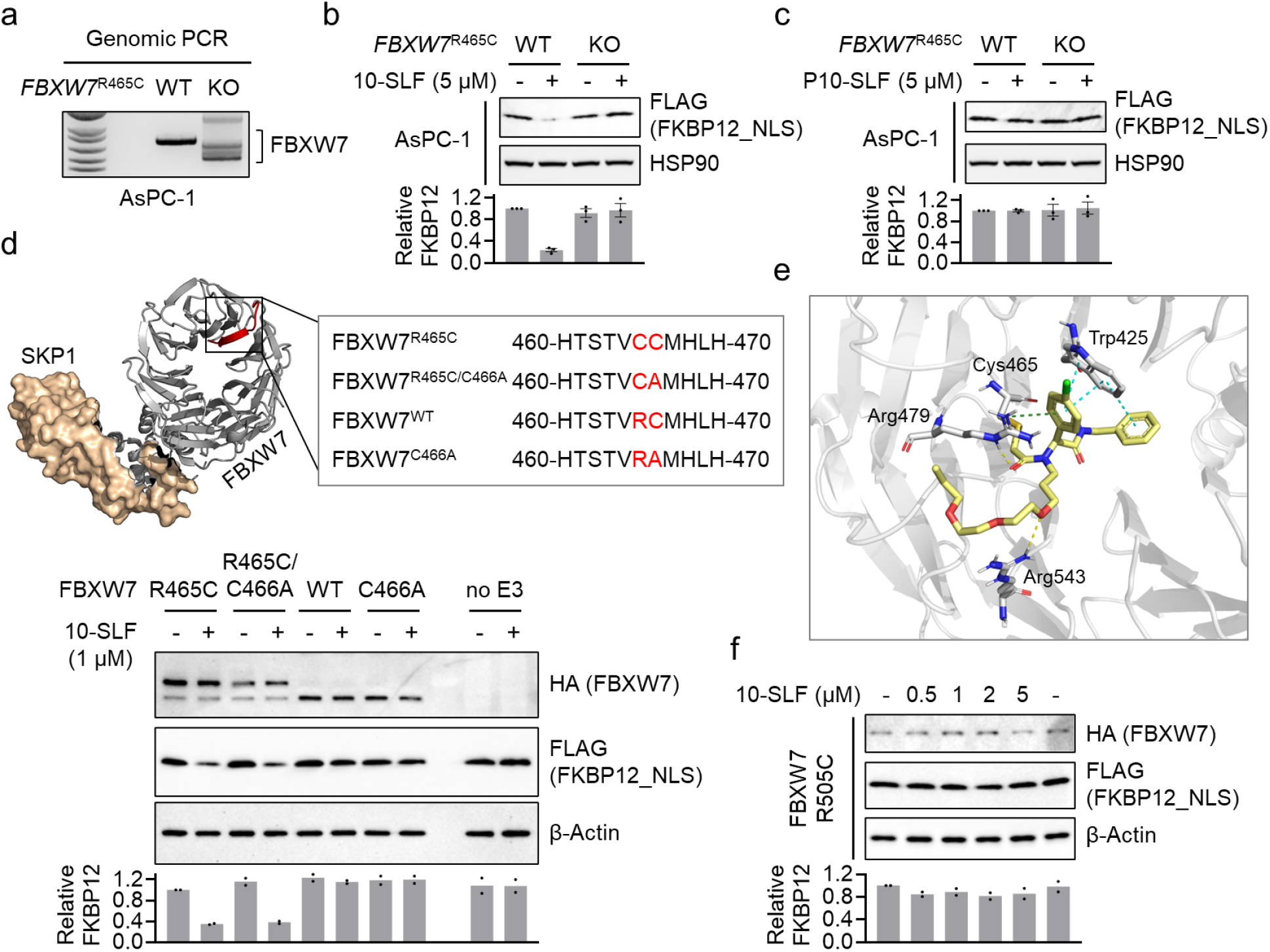
Mechanistic insights into FBXW7-R465C-mediated target degradation. **a**. Genomic PCR confirms *FKBW7*^R465^ KO in AsPC-1 cells. **b**. 10-SLF induced FKBP12_NLS degradation in AsPC-1 WT but not *FKBW7*^R465^ KO cells. The bar graph represents quantification of the FKBP12/HSP90 protein content. Data are presented as the mean values ± s.e.m. (n = 3 independent replicates). **c**. P10-SLF did not induce FKBP12_NLS degradation in either AsPC-1 WT or *FKBW7*^R465^ KO cells. The bar graph represents quantification of the FKBP12/HSP90 protein content. Data are presented as the mean values ± s.e.m. (n = 3 independent replicates). **d**. A series of mutations were introduced in FBXW7 for this study. The crystal structure was obtained from the Protein Data Bank (PDB: 7T1Y). The bottom panel shows the evaluation of different FBXW7 mutations in supporting 10-SLF-induced FKBP12 degradation. The bar graph represents quantification of the FKBP12/*β*-Actin protein content. Data are presented as the mean values (n = 2 independent replicates). **e**. The modeling study revealed interactions between the electrophilic portion of 10-SLF and a pocket in FBXW7 involving R465C. **f**. FBXW7-R505C did not support the degradation of FKBP12_NLS induced by 10-SLF. The bar graph represents quantification of the FKBP12/*β*-Actin protein content. Data are presented as the mean values (n = 2 independent replicates).

FBXW7-R465C contains two consecutive cysteines (C465 and C466). Since FBXW7 WT does not support 10-SLF-induced FKBP12 degradation, we hypothesize that C466 itself does not serve as a modified site for inducing target degradation by 10-SLF. Nonetheless, we wondered whether C466 might contribute to degradation activity in the context of FBXW7-R465C. To investigate this, we generated a series of mutations in FBXW7, including R465C, R465C/C466A, and C466A (**Figure 3d**), and measured their ability to support 10-SLF-induced FKBP12 degradation. The results indicated that FBXW7-R465C and FBXW7-R465C/C466A had similar activities in inducing FKBP12 degradation by 10-SLF (**Figure 3d**), suggesting a non-essential role for C466 in this process. To gain further insights into the binding model between 10-SLF and FBXW7-R465C, we conducted a docking study using the FBXW7 binding portion of 10-SLF and a crystal structure of FBXW7 (PDB: 7T1Y). The results indicated a favorable fit of 10-SLF on the surface of FBXW7 incorporating the manually introduced R465C mutation. The α-chloroacetamide formed a covalent bond with C465, and its carbonyl group formed a hydrogen bond with R479. Additionally, a hydrogen bond was observed between the oxygen atom of the polyethylene glycol linker and R543. The 4-chlorophenyl group formed a cation–π interaction with R479 and a π–π interaction with W425 (**Figure 3e**). We further investigated another hotspot mutation, R505C, located near R465C (**Figure S3**). Our results revealed no FKBP12 degradation by 10-SLF when FBXW7-R505C was expressed (**Figure 3f**). While identifying ligands that engage FBXW7-R505C for TPD applications represents a promising avenue for future research, our initial compound 10-SLF appears to rely on FBXW7-R465C, rather than R505C, for target degradation.

Next, we aimed to measure the formation of a ternary complex involving FKBP12, FBXW7-R465C, and 10-SLF, a critical step previously shown to drive target degradation^23^. HEK293T cells expressing FLAG-FKBP12_NLS and HA-FBXW7 variants were treated with 10-SLF and MG132, followed by cell lysis and FKBP12 immunoprecipitation. The results indicated that both FBXW7-R465C and FBXW7-R465C/C466A coimmunoprecipitated with FKBP12 in the presence of 10-SLF (**Figure 4a**), supporting the formation of a ternary complex with these mutants and the essential role of R465C in this process. In contrast, the formation of the ternary complex was significantly hindered when FBXW7 WT or FBXW7-C466A was expressed (**Figure 4a**).

**Figure 4.**
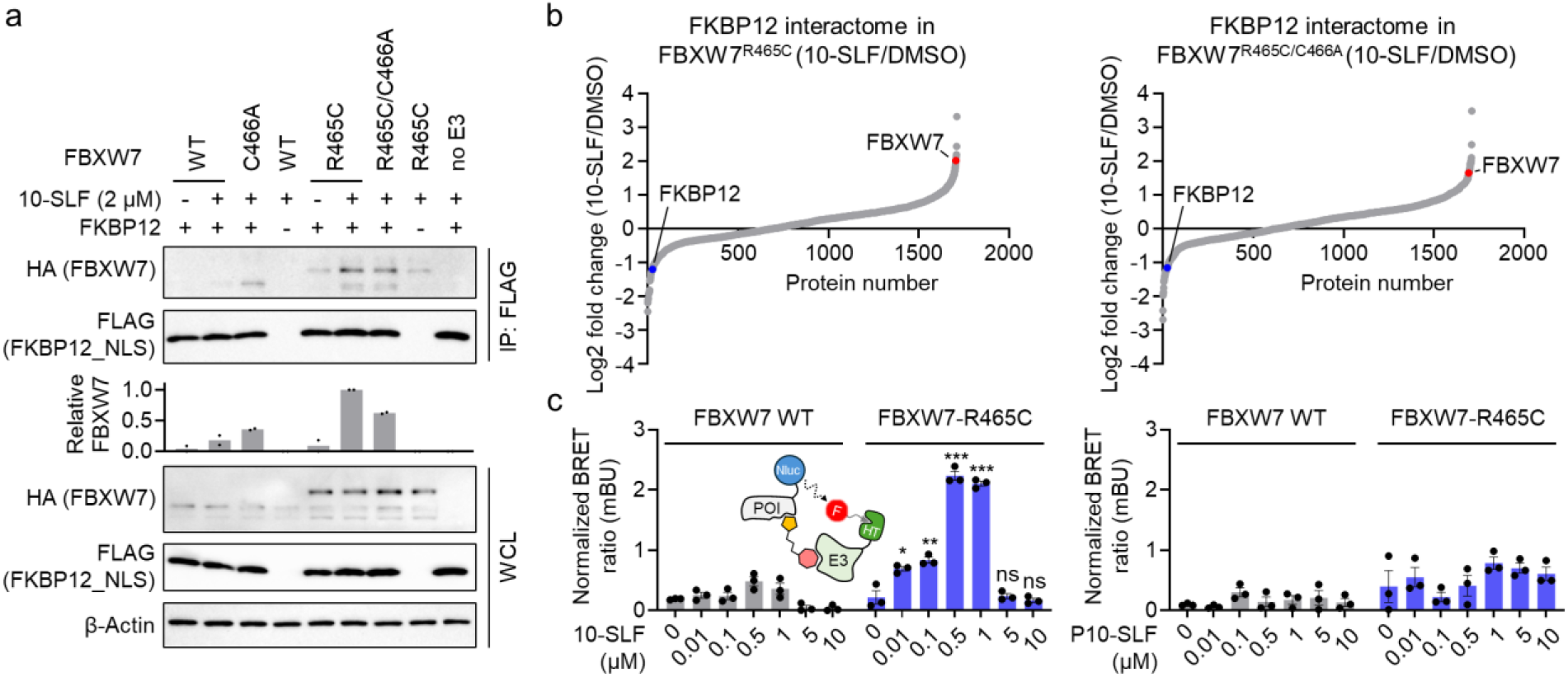
10-SLF induces the formation of a ternary complex between FKBP12 and FBXW7-R465C. **a**. Coimmunoprecipitation assays demonstrated that HA-FBXW7-R465C and HA-FBXW7-R465C/C466A coimmunoprecipitated with FLAG-FKBP12_NLS in the presence of 10-SLF and MG132. The bar graph represents quantification of the immunoprecipitated HA-FBXW7 compared to HA-FBXW7 in whole cell lysates (WCL). Data are presented as the mean values (n = 2 independent replicates). **b**. FKBP12 enrichment proteomic analysis revealed that HA-FBXW7-R465C and HA-FBXW7-R465C/C466A interacted with FLAG-FKBP12_NLS in cells treated with 10-SLF. **c**. A NanoBRET assay measuring the interaction between FKBP12 and FBXW7 indicated that 10-SLF induced this interaction with FBXW7-R465C, but not with FBXW7 WT. Data are presented as the mean values ± s.e.m. (n = 3 independent replicates). The statistical significance was evaluated through unpaired two-tailed Student’s *t*-tests, comparing cells treated with 10-SLF to DMSO. \**P* < 0.05, \*\**P* < 0.01, \*\*\**P* < 0.001, and ns: not significant.

To further explore the FKBP12 interactome landscape affected by 10-SLF, we employed affinity purification-mass spectrometry (AP-MS) to identify proteins interacting with FLAG-FKBP12_NLS from HEK293T cells treated with 10-SLF. The results showed that FBXW7-R465C and FBXW7-R465C/C466A were recruited by 10-SLF (**Figure 4b** and **Table S2**). Additionally, we established a NanoBRET assay to directly measure the interactions between FKBP12 and FBXW7 (**Figure 4c**). The results indicated that only in cells expressing FBXW7-R465C did 10-SLF induce the interaction between FKBP12 and FBXW7, whereas P10-SLF did not (**Figure 4c**). Notably, we observed a hook effect in the NanoBRET assay at concentrations of 5 and 10 µM of 10-SLF (**Figure 4c**). Collectively, these findings indicate that 10-SLF promotes the formation of a ternary complex involving FKBP12, FBXW7-R465C, and 10-SLF.

Finally, we aimed to assess the potential of harnessing FBXW7-R465C to facilitate the degradation of additional protein targets. We synthesized a heterobifunctional compound, 10-MKI, by coupling the FBXW7-R465C binding moiety to a multi-kinase binder previously used to explore degradable kinases^24^ (**Figure 5a**). We conducted a global proteomic analysis in AsPC-1 parental and *FBXW7*^R465C^ KO cells treated with 10-MKI, identifying a handful of kinases that were more degraded in the parental cells compared to the *FBXW7*^R465C^ KO cells (**Figure 5b** and **Table S3**). We selected AURKA as a candidate kinase and validated its degradation by 10-MKI through Western blot analysis in AsPC-1 parental cells, while no degradation was observed in *FBXW7*^R465C^ KO cells (**Figure 5c**). Additionally, we tested three other cell lines: CCRF-CEM (expressing *FBXW7*^R465C^), HEK293T (expressing *FBXW7* WT), and A549 (expressing *FBXW7* WT). Our results showed that 10-MKI degraded AURKA and CSNK1D only in the CCRF-CEM cell line expressing *FBXW7*^R465C^, while no degradation occurred in the other two cell lines expressing *FBXW7* WT (**Figure 5d**). We further employed the NanoBRET assays to measure the formation of a ternary complex between AURKA and FBXW7-R465C induced by 10-MKI, along with the ubiquitination of FBXW7-R465C and the proteasomal recruitment of AURKA. D1 (**Figure S4)**, a CRBN-based multi-kinase degrader^22^, served as a positive control. The results indicated that in cells expressing FBXW7-R465C, 10-MKI effectively promoted ternary complex formation, AURKA ubiquitination, and proteasomal recruitment (**Figure 5e**). These induced actions were completely blocked by the E1 enzyme inhibitor, further confirming the involvement of the proteasome-mediated degradation mechanism. Notably, despite the interaction between AURKA and FBXW7 WT induced by 10-MKI at higher concentrations (5 and 10 µM) (**Figure 5e**), no ubiquitination or degradation of AURKA was observed under these conditions.

**Figure 5.**
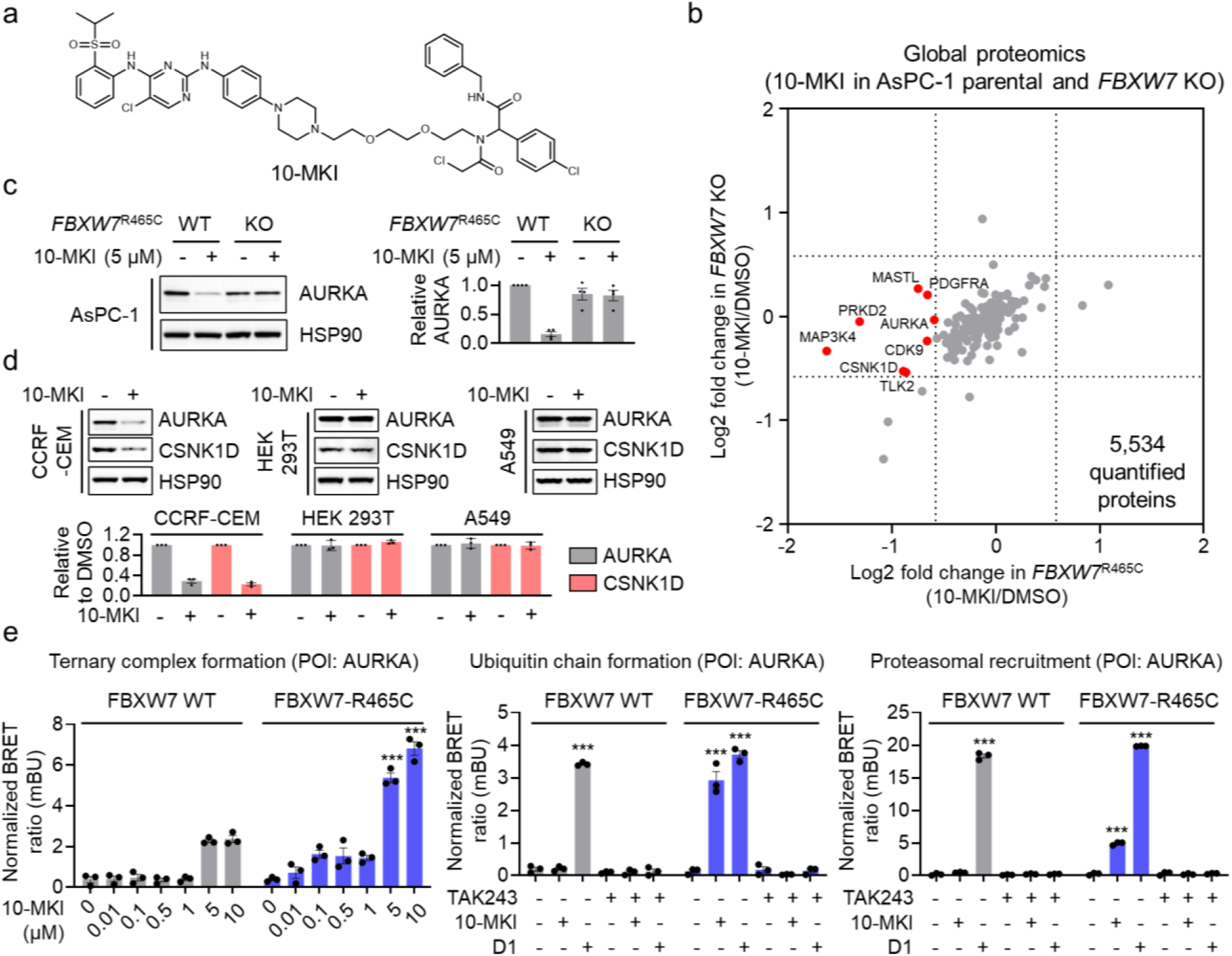
Utilizing FBXW7-R465C for the degradation of additional protein targets. **a**. Structure of 10-MKI. **b**. Global proteomic analysis in AsPC-1 parental and *FBXW7*^R465C^ KO cells treated with 5 μM 10-MKI for 24 hours. Data are presented as the mean values (n = 2 independent replicates for DMSO treatment, n = 3 independent replicates for 10-MKI treatment). **c**. Treatment with 10-MKI (5 μM, 24 hours) induced AURKA degradation in AsPC-1 parental cells but not in *FBXW7*^R465C^ KO cells. The bar graph represents quantification of the AURKA/HSP90 protein content. Data are presented as the mean values ± s.e.m. (n = 3 independent replicates). **d**. 10-MKI induced the degradation of AURKA and CSNK1D in CCRF-CEM cells, but not in HEK293T or A549 cells. The bar graph represents quantification of the AURKA/HSP90 protein content. Data are presented as the mean values ± s.e.m. (n = 3 independent replicates). **e**. NanoBRET assays demonstrated that 10-MKI induced ternary complex formation between FBXW7-R465C and AURKA, along with the ubiquitination and proteasomal engagement of AURKA. Data are presented as the mean values ± s.e.m. (n = 3 independent replicates). The statistical significance was evaluated through unpaired two-tailed Student’s *t*-tests, comparing cells treated with 10-MKI or D1 to DMSO. \*\*\**P* < 0.001.

In summary, this study demonstrates that the FBXW7 somatic mutation R465C can be leveraged by heterobifunctional compounds to degrade multiple target proteins. This discovery underscores the potential of harnessing mutant E3 ligases specific to diseased cells for TPD applications.

## Acknowledgement

We gratefully acknowledge the support of the NIH R00 CA248715 (X.Z.), NIH T32 GM105538 (A.A.B.), Damon Runyon Cancer Research Foundation DFS-53-22 (X.Z.), and the Chemistry of Life Processes Institute Cornew Innovation Award (X.Z.).

## Notes

The authors declare no competing financial interest.

