## Supplementary Information for "Harnessing the FBXW7 somatic mutant R465C for targeted protein degradation"

### Supplementary Materials and Methods

#### Reagents

The anti-HA (clone#: C29F4, cat#: 3724, dilution 1:5000), HRP-linked anti-HSP90 (clone#: C45G5, cat#: 79641, dilution 1:5000), anti-AURKA (clone#: D3E4Q, cat#: 14475, dilution 1:1000), HRP-linked rabbit IgG (cat#: 7074, dilution 1:5000), and anti-CSNK1D antibody (polyclonal, cat#: 12417, dilution 1:1000) antibodies were purchased from Cell Signaling Technology. The anti-FLAG HRP antibody (clone M2, cat#: A8592, dilution 1:5000), anti-FLAG affinity gel (clone M2, cat#: A2220), anti-HA HRP antibody (clone#: 3F10, cat#: 12013819001, dilution 1:5000), and cOmplete protease inhibitor cocktail (cat#: 11873580001) were purchased from Sigma-Aldrich. The anti-β-Actin HRP antibody (clone#: C4, cat#: sc-47778, dilution 1:1000) was purchased from Santa Cruz Biotechnology. Cas9 endonuclease was purchased from Integrated DNA Technologies (cat# 1081061). PureLink genomic DNA mini kit (cat#: K182001) was purchased from Invitrogen. FuGene 6 (cat#: E2692) transfection reagent and sequencing grade trypsin/Lys-C mix (cat#: V5071) were purchased from Promega. Puromycin (cat#: ant-pr-1) and blastcidin (cat#: ant-bl-05) were purchased from InvivoGen. MG132 (cat#: S2619) was purchased from Selleck Chemicals. MLN4924 (cat#: 15217) and SLF (cat#: 10007974) were purchased from Cayman Chemical. Enzyme-linked chemiluminescence (ECL) (cat#: 32106), ECL plus (cat#: 32132), BCA protein assay kit (cat#: 23227), and Tandem Mass Tag (TMT) isobaric label reagent (cat#: 90110) were purchased from Thermo Scientific.

#### Cell Lines

HEK293T and A549 cells were obtained from ATCC. AsPC-1 and CCRF-CEM cells were obtained from The Developmental Therapeutics Core at Northwestern University. HEK293T cells were cultured in Dulbecco's Modified Eagle Medium (DMEM, Corning, cat# 15013CV) with 10% (v/v) fetal bovine serum (FBS, Omega Scientific, cat# FB-01) and L-glutamine (2mM, Gibco, cat# 25030081). A549 cells were cultured in DMEM with 10% (v/v) FBS, L-glutamine (2mM) and MEM non-essential amino acids (Gibco, cat# 11140050). AsPC-1 and CCRF-CEM cells were cultured in RPMI-1640 medium (Corning, cat# 15040CV) with 10% (v/v) FBS and L-glutamine (2mM). All the cell lines were tested negative for mycoplasma contamination using Universal Mycoplasma Detection Kit (ATCC, cat# 30-1012K).

#### Generation of *FBXW7* Knockout Cells

HEK293T and AsPC-1 cells with *FBXW7* CRISPR-Cas9 knockout were generated by electroporating a complex consisting of Cas9 and three sgRNAs using the 4D-Nucleofector (Lonza Bioscience). Three sgRNAs targeting *FBXW7* gene are: *FBXW7* sgRNA#1: UCGGUAGAUGAGGACUCCUC; *FBXW7* sgRNA#2: CACGAACUCCAGUAGUAUUG; *FBXW7* sgRNA#3: GUUGGAGUAGAACCUAGACC. To confirm the knockout of *FBXW7* gene, genomic DNA was extracted using the PureLink Genomic DNA Mini Kit *FBXW7* gene was then amplified by PCR and measured through DNA gel electrophoresis and DNA sequencing. The sequencing primers used for *FBXW7* were: 5'-TAGCCAAGGTCCAAGAAGTAGC-3' (forward) and 5'-CCTCCATTTGTACTCAGATTGTCC-3' (reverse).

#### Cloning, Mutagenesis, and Lentivirus Transduction

Human *FBXW7* with an N-terminal HA tag was purchased as a gene block from Integrated DNA Technologies and cloned into pCDH-CMV-MCS-EF1-Blast vector. The FLAG-FKBP12_NLS construct was generated as previously described[1]. FBXW7 mutants were created using the Q5 Site-Directed Mutagenesis Kit (New England Biolabs). Lentivirus was generated by co-transfecting HEK293T cells with a plasmid containing the gene of interest, psPAX2, and pMD2.G using FuGene 6 transfection reagent. The medium containing lentiviral particles was collected after 48 hours, filtered through a 0.45 μm Millex-HV sterile syringe filter unit (MilliporeSigma), and used to transduce HEK293T or AsPC-1 cells. To generate stable cell lines, puromycin (2 μg/mL) or blasticidin (10 μg/mL) was added and incubated with the cells for 7 days.

#### Cell Lysis and Western Blot

Cells were lysed in radioimmunoprecipitation assay (RIPA) buffer containing 25 mM Tris-HCl (pH 7.6), 150 mM NaCl, 1% NP-40, 1% sodium deoxycholate, and 0.1% SDS. The lysis buffer was supplemented with the cOmplete protease inhibitor cocktail prior to use. The protein concentration was quantified using the BCA assay. The protein lysate was mixed with Laemmli sample buffer and heated at 95°C for 5 minutes. Proteins were separated using 4-20% Novex Tris-Glycine gels (Thermo Scientific, cat# XP04205BOX) and transferred onto a 0.2 µm polyvinylidene fluoride (PVDF) membrane (Bio-Rad, cat# 1620177). The PVDF membrane was blocked with a solution of 5% non-fat milk in TBST buffer (0.1% Tween 20, 20 mM Tris-HCl at pH 7.6, and 150 mM NaCl) for 1 hour at room temperature. Primary antibodies, diluted in 5% non-fat milk in TBST buffer, were applied to the membrane. Incubation times were 1 hour at room temperature for FLAG, HA, and β-Actin antibodies, and overnight at 4°C for the other antibodies. After primary antibody incubation, the membrane was washed three times with TBST buffer and then incubated with a secondary antibody for 1 hour at room temperature. Following three additional washes with TBST buffer, the chemiluminescence signal on the membrane was developed using ECL western blotting detection reagent, and the resulting signal was measured with the ChemiDoc MP system (Bio-Rad). Band intensities were quantified using ImageJ software (version 1.51h).

**Immunoprecipitations**

Cells were lysed in NP-40 lysis buffer (25 mM Tris-HCl, pH 7.4; 150 mM NaCl; 10% glycerol; 1% NP-40; cOmplete protease inhibitor cocktail). The lysate was centrifuged at 16,000 g for 10 minutes at 4°C, and the supernatant was collected for immunoprecipitation. FLAG gel (25 µL slurry per sample) was added to the protein lysates and rotated at 4°C for 2 hours. The affinity gel was then washed four times with immunoprecipitation washing buffer (25 mM Tris-HCl, pH 7.4; 150 mM NaCl; 0.2% NP-40). The washed affinity gel was mixed with Laemmli sample buffer and heated at 95°C for 10 minutes. The resulting supernatant was collected for western blot analysis.

**NanoBRET Ubiquitination, Ternary Complex Formation (TCF), and Proteasomal Recruitment Assays**

To assess the formation of FKBP12-FBXW7, FKBP12-FBXW7-R465C, AURKA-FBXW7 or AURKA-FBXW7-R465C complexes, HEK293T cells ($1 \times 10^6$) were maintained in complete medium and transfected with 20 ng Nluc-FKBP12- or Nluc-AURKA, and 2 µg of either HaloTag-FBXW7, or HaloTag-FBXW7-R465C vectors in six-well plates using FuGENE HD (Promega). To directly evaluate live-cell FKBP12 or AURKA ubiquitination and FKBP12 or AURKA proteasomal recruitment, *FBXW7* KO HEK293T cells or *FBXW7*

KO HEK293T cells stably expressing FBXW7-R465C ($1 \times 10^6$) were transfected with 20 ng Nluc-FKBP12- or Nluc-AURKA, and 2 µg of either HaloTag-Ubiquitin, or HaloTag-PSMD3 vectors in six-well plates using FuGENE HD. The following day, transfected cells ($2 \times 10^4$) were replated in triplicate into white 96-well tissue culture plates in 90 µL of assay medium (Opti-MEM Reduced Serum Medium, no phenol red, and 4% FBS) in the presence or absence of HaloTag NanoBRET 618 Ligand (Promega) and incubated overnight at 37°C and 5% $CO_2$. To perform end-point monitoring of ternary complex formation, ubiquitination, and proteasomal recruitment, 10 µL of 10X concentrated compounds were added to the related wells and incubated at 37°C, 5% $CO_2$ for 7 hours. Then, 25 µL of 5X NanoBRET Nano-Glo (Promega) substrate was added, and BRET was measured using VICTOR Nivo Multimode Plate Reader (PerkinElmer, USA). Dual-filtered luminescence was collected with a 460/480-nm bandpass filter (donor, FKBP12- or AURKA-NanoLuc fusion proteins) and a 610-nm long pass filter (acceptor, HaloTag NanoBRET 618 ligand) using an integration time of 0.5 s. For all NanoBRET experiments, background subtracted NanoBRET ratios were expressed in milliBRET units.

**Global proteomics**

Cells were lysed in 100 µL of PBS by sonication (10 pulses at 40% intensity, three rounds). Protein concentration was determined using a DC assay (Bio-Rad, cat# 5000112). A total of 100 µg of protein in 100 µL of lysis buffer was denatured with 8 M urea. For reduction, 5 µL of a 200 mM DTT solution in water was added, and the sample was heated to 65°C for 15 minutes. Alkylation was performed by adding 5 µL of a 400 mM iodoacetamide solution in water, followed by incubation in the dark at 37°C for 30 minutes. Proteins were then precipitated by adding 600 µL of MeOH, 200 µL of $CHC_{l3}$, and 500 µL of water. The resulting protein pellets were washed with 1 mL of MeOH, then solubilized in 160 µL of 200 mM EPPS buffer. Each sample was digested with 2 µg of LysC enzyme at 37°C for 2 hours, followed by the addition of 5 µg of trypsin/LysC mix for further digestion at 37°C for 12 hours. For TMT labeling, 12.5 µg of peptides in 35 µL of EPPS buffer were prepared, to which 9 µL of $CH_3CN$ and TMT tags (3 µL per sample, 20 µg/µL in $CH_3CN$) were added. Samples were incubated at room temperature for 1 hour, and the reaction was quenched by adding 6 µL of a 5% hydroxylamine solution and 2.5 µL of formic acid. Labeled samples were pooled and separated into 12 fractions using a Thermo Vanquish UHPLC fractionator. Peptide fractions were analyzed by liquid chromatography-tandem mass spectrometry on an Orbitrap Eclipse Tribrid Mass Spectrometer coupled with a Vanquish

Neo UHPLC system. Peptides were loaded onto an EASY-Spray HPLC column (C18, 2 µm particle size, 75 µm inner diameter, 250 mm length) and eluted at a flow rate of 0.25 µL/min using the following gradient: 5% buffer B (80% acetonitrile with 0.1% formic acid) in buffer A (water with 0.1% formic acid) from 0 to 15 minutes, increasing to 45% buffer B from 15 to 155 minutes, and then from 45% to 100% buffer B from 155 to 180 minutes. The nano-LC electrospray ionization source was set at 1.5 kV. The analysis began with an MS1 scan (Orbitrap, 120,000 resolution, m/z range 375–1600, RF lens 30%, standard AGC target, auto maximum injection time). In MS2, precursor ions were isolated by quadrupole (0.7 isolation window) and fragmented via HCD in the ion trap (collision energy 32%, standard AGC, maximum injection time 35 ms). After each MS2 scan, synchronous precursor selection (SPS) enabled selection of up to 10 MS2 fragment ions for MS3 analysis, where precursors were fragmented by HCD and analyzed in the Orbitrap (collision energy 55%, AGC target 250%, maximum injection time 200 ms, 50,000 resolution). RAW data were collected in Xcalibur (version 4.5.445.18) and analyzed in Proteome Discoverer 2.5.

**Affinity Purification–Mass Spectrometry**

Cells were lysed in NP-40 lysis buffer with the cOmplete protease inhibitor cocktail. After centrifugation at 16,000 g for 10 minutes at 4°C, the supernatant was collected for immunoprecipitation. Protein lysates were incubated with 25 µL FLAG affinity gel slurry per sample for 2 hours at 4°C. The gel was washed four times in an immunoprecipitation washing buffer, followed by two washes with PBS. FLAG-FKBP12 and the associated proteins were eluted by heating the beads at 65°C for 10 minutes in 8 M urea in PBS. The eluted proteins were then reduced with 12.5 mM DTT at 65°C for 15 minutes, followed by alkylation with 25 mM iodoacetamide at 37°C for 30 minutes. The protein solution was diluted in PBS to achieve a urea concentration of 2 M and digested with 2 µg of trypsin/Lys-C mix at 37°C for 12 hours. TMT tags (6 µL, 20 µg/µL in dry $CH_3CN$) were added and incubated at room temperature for 1 hour, after which the reaction was quenched with 6 µL of a 5% hydroxylamine solution and 2.5 µL of formic acid. Samples were then pooled and desalted using a Sep-Pak C18 cartridge (Waters, cat# WAT054955). The eluted peptide solution was dried with a SpeedVac concentrator and analyzed by LC-MS using the methodology described above.

### Modeling Study

The crystal structure of FBXW7 from the Protein Data Bank (PDB: 7T1Y) was used for covalent docking. Protein preparation was conducted in Maestro 13.4 (Schrödinger) to preprocess, optimize hydrogen bond assignment, perform energy minimization, and remove water molecules. The compound was prepared with the LigPrep module using the OPLS4 force field. Docking was carried out using the CovDock module, with nucleophilic substitution selected as the reaction type and default parameters applied for other settings. The final pose was chosen based on low-energy conformation and favorable hydrogen-bond, cation-pi, and pi-pi interaction geometries. Figures were generated using PyMOL.

### Statistical Analysis

Quantitative data are presented as scatter plots, with the mean displayed alongside standard error of the mean (s.e.m.) as error bars. Comparisons between two groups were analyzed using an unpaired two-tailed Student's *t*-test. Significance levels are indicated as follows: *$P < 0.05$, **$P < 0.01$, and ***$P < 0.001$.

### Synthesis of P10-SLF

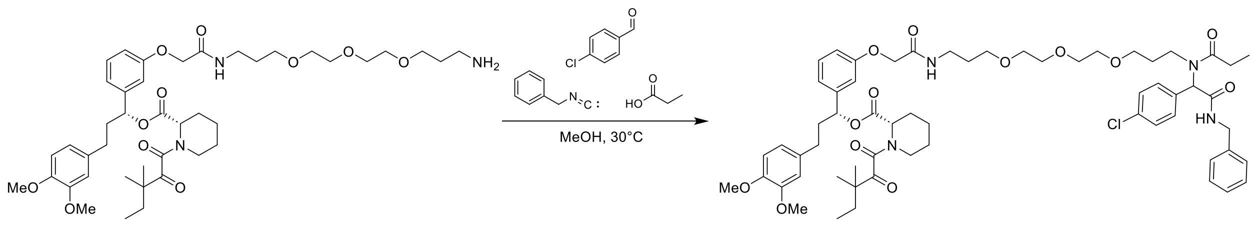


S-1 P10-SLF

S-1 was prepared as reported previously[2]. S-1 (200 mg, 0.25 mmol, 1 eq) and p-chlorobenzaldehyde (36 mg, 0.25 mmol, 1.0 eq) was stirred in MeOH (1.0 mL) at 30℃ for 12 h. After this, the benzyl isocyanide (35 mg, 0.30 mmol, 1.2 eq) and propionic acid (22 mg, 0.30 mmol, 1.2 eq) were added and the reaction was stirred for 24 h. MeOH was removed and the residue was purified by preparative TLC to provide the desired analogue P10-SLF (140 mg, 0.13 mmol, 52%).

**¹H NMR** (500 MHz, $CDCl_3$) δ ppm 7.33-7.29 (m, 7H), 7.23-7.20 (m, 3H), 7.01-6.94 (m, 2H), 6.91 (s, 1H), 6.84-6.82 (m, 1H), 6.78-6.75 (m, 1H), 6.69-6.63 (m, 3H), 5.77-5.73 (m, 2H), 4.45-4.42 (m, 5H), 3.84-3.83 (m, 6H), 3.57-3.55 (m, 2H), 3.54-3.50 (m, 6H), 3.45-3.33 (m, 8H), 3.27-3.25 (m, 2H), 3.18-3.12 (m, 1H), 2.46-2.40 (m, 2H), 2.25-2.20 (m, 1H), 2.05-2.02 (m, 1H), 1.83-1.77 (m, 3H), 1.75-1.60 (m, 6H), 1.50-1.45 (m, 1H), 1.41-1.31 (m, 2H), 1.20 (s, 3H), 1.89 (s, 3H), 1.14-1.10 (m, 4H), 0.87 (t, $J$ = 7.5 Hz, 3H).

**¹³C NMR** (126 MHz, $CDCl_3$) δ 207.92, 175.15, 169.73, 168.02, 167.36, 157.59, 148.99, 147.47, 141.94, 138.10, 134.36, 134.15, 133.42, 130.76, 130.08, 128.99, 128.69, 127.71, 127.46, 120.23, 120.15, 114.33, 113.38, 113.37, 111.82, 111.43, 111.41, 76.55, 70.62, 70.55, 70.34, 70.26, 69.54, 69.52, 68.20, 67.53, 62.52, 56.01, 55.92, 51.33, 46.77, 44.52, 44.23, 43.68, 41.07, 38.27, 37.18, 32.55, 31.33, 29.82, 29.36, 26.68, 26.50, 25.02, 23.56, 23.54, 23.24, 21.28, 9.61, 8.84.

**HRMS** (ESI+) *m/z* calcd for $C_{60}H_{79}ClN_4NaO_{13}^+$ $[M+Na]^+$: 1121.5225, found 1121.5259.

¹H NMR spectrum of P10-SLF

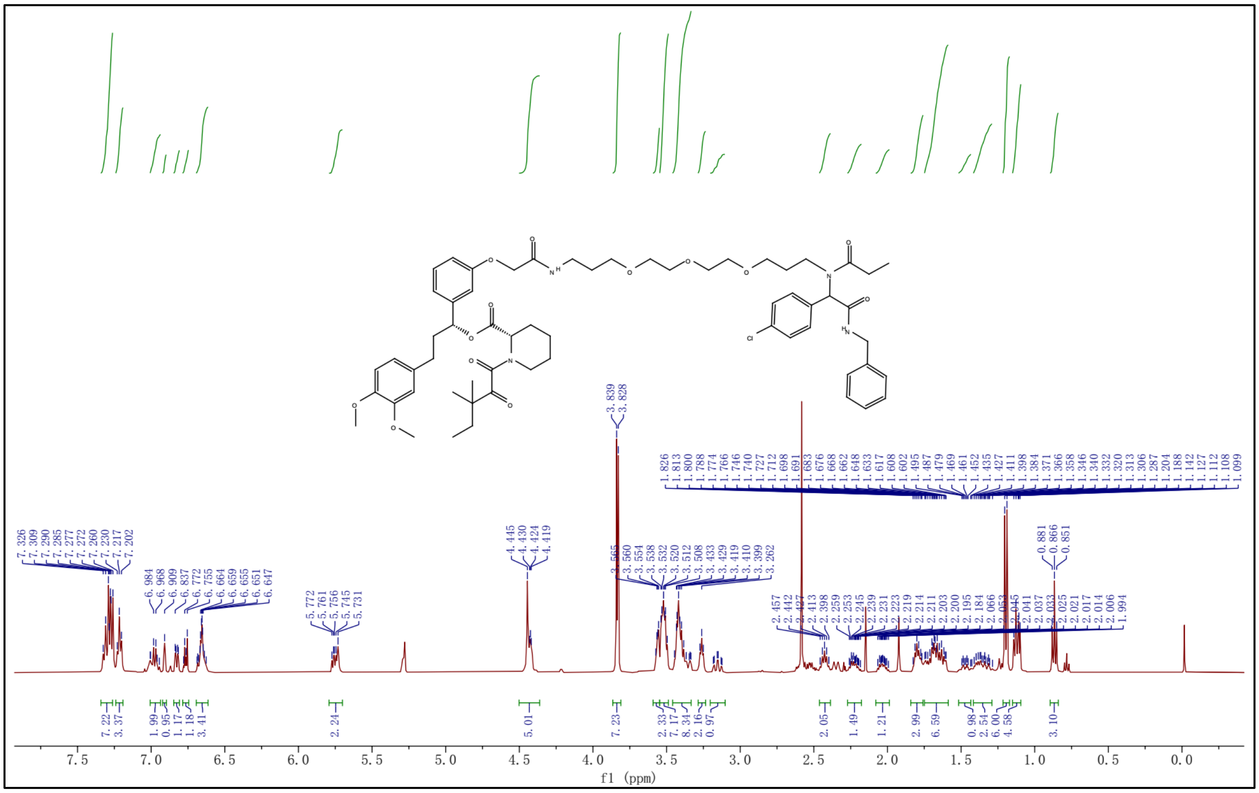

$^{13}C$ NMR spectrum of P10-SLF

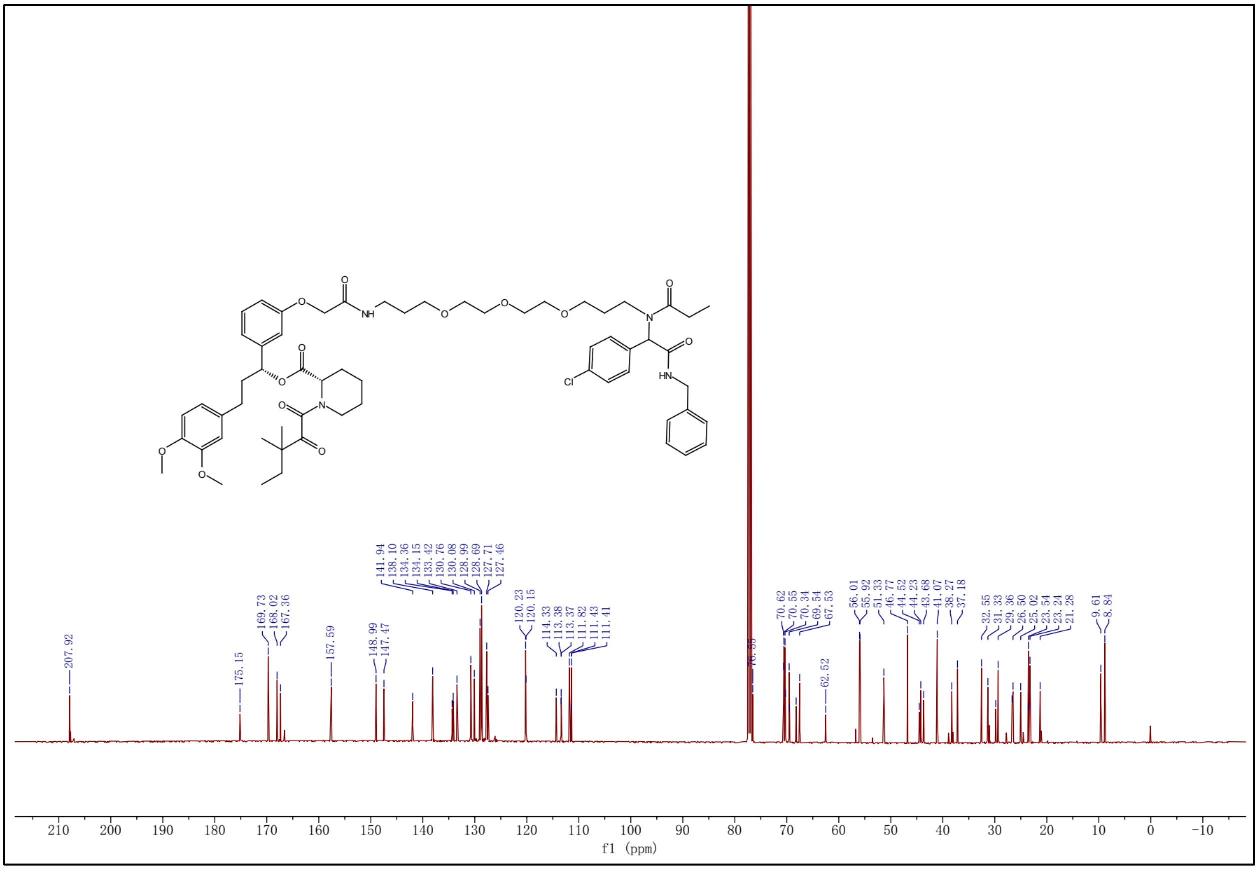


### Synthesis of 10-MKI

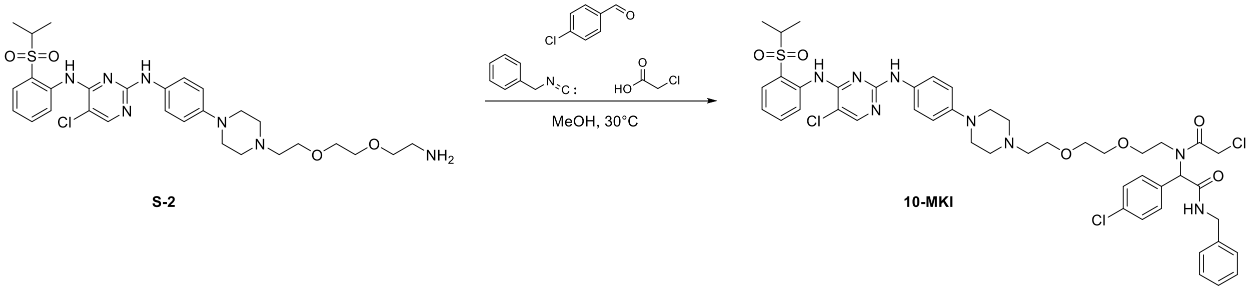


S-2 was prepared as reported previously[3]. S-2 (150 mg, 0.24 mmol, 1 eq) and p-chlorobenzaldehyde (34 mg, 0.24 mmol, 1.0 eq) was stirred in MeOH (1.0 mL, 0.5 M) at 30℃ for 24 h. After this, the benzyl isocyanide (34 mg, 0.29 mmol, 1.2 eq) and chloroacetic acid (27 mg, 0.29 mmol, 1.2 eq) were added and the reaction was stirred for 12 h. MeOH was removed and the residue was purified by preparative TLC to provide the desired analogue 10-MKI (105 mg, 0.11 mmol, 46%).

**$^1$H NMR** (500 MHz, $CDCl_3$) δ ppm 9.61 (s, 1H), 8.58 (d, *J* = 10.0 Hz, 1H), 8.10 (s, 1H), 7.89 (d, *J* = 10.0 Hz, 1H), 7.56 (t, 1H, *J* = 5.0 Hz), 7.38 (d, 2H, *J* = 10.0 Hz), 7.34-7.32 (m, 4H), 7.30 (d, 2H, *J* = 5.0 Hz), 7.24-7.21 (m, 2H), 6.93 (s, 1H), 6.89 (d, 2H, *J* = 10.0 Hz), 6.68 (s, 1H), 5.81 (s, 1H), 4.48-4.39 (m, 4H), 3.66-3.60 (m, 3H), 3.51-3.40 (m, 6H), 3.26-3.18 (m, 6H), 2.80-2.74 (m, 6H), 1.31 (d, *J* = 5.0 Hz, 6H).

**$^{13}$C NMR** (126 MHz, $CDCl_3$) δ 169.01, 168.93, 158.22, 155.51, 155.31, 138.60, 137.94, 134.90, 134.63, 133.25, 132.00, 131.36, 130.84, 129.31, 128.85, 127.86, 127.68, 124.51, 123.58, 123.22, 122.23, 117.08, 105.97, 70.61, 70.31, 69.09, 68.39, 63.03, 57.70, 55.74, 53.58, 49.37, 46.80, 43.91, 42.31, 15.50.

**HRMS** (ESI+) *m/z* calcd for $C_{46}H_{54}Cl_3N_8O_6S^+$ $[M+H]^+$: 953.2918, found 953.2999.

$^1$H NMR spectrum of 10-MKI

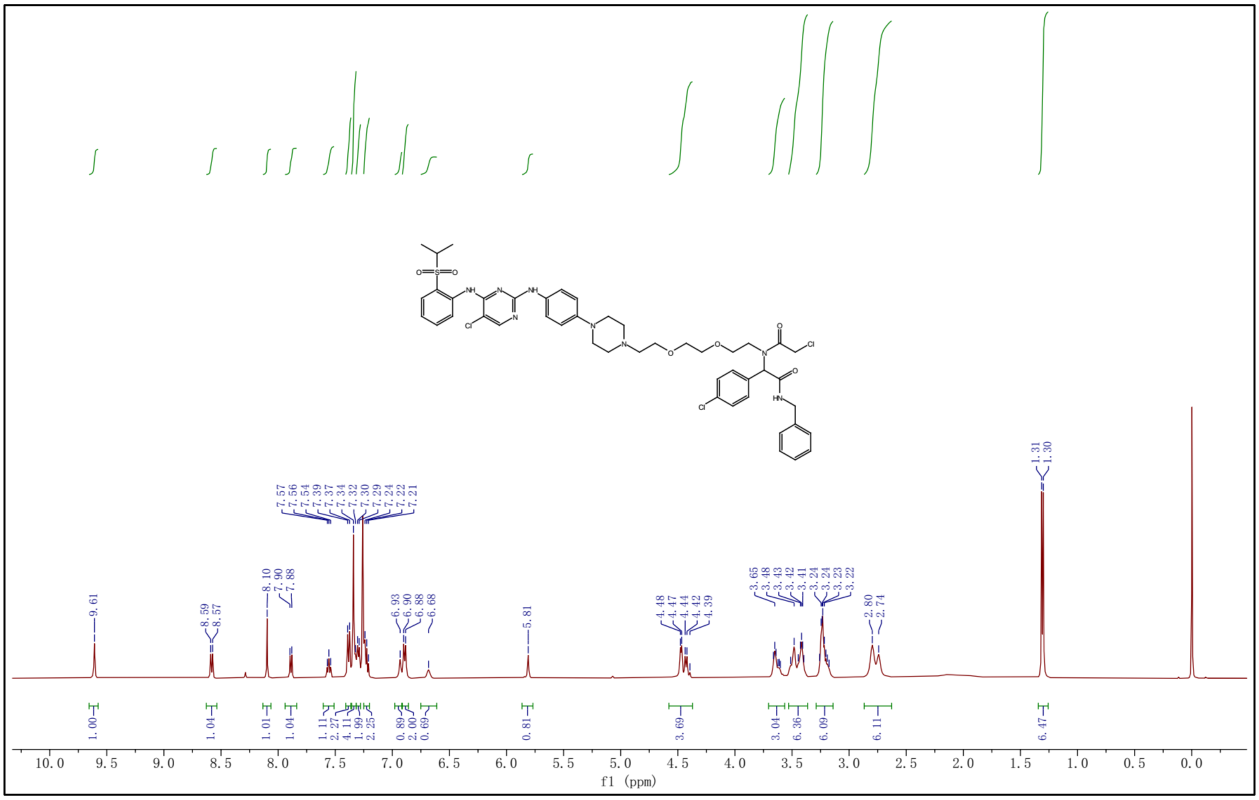

$^{13}C$ NMR spectrum of 10-MKI

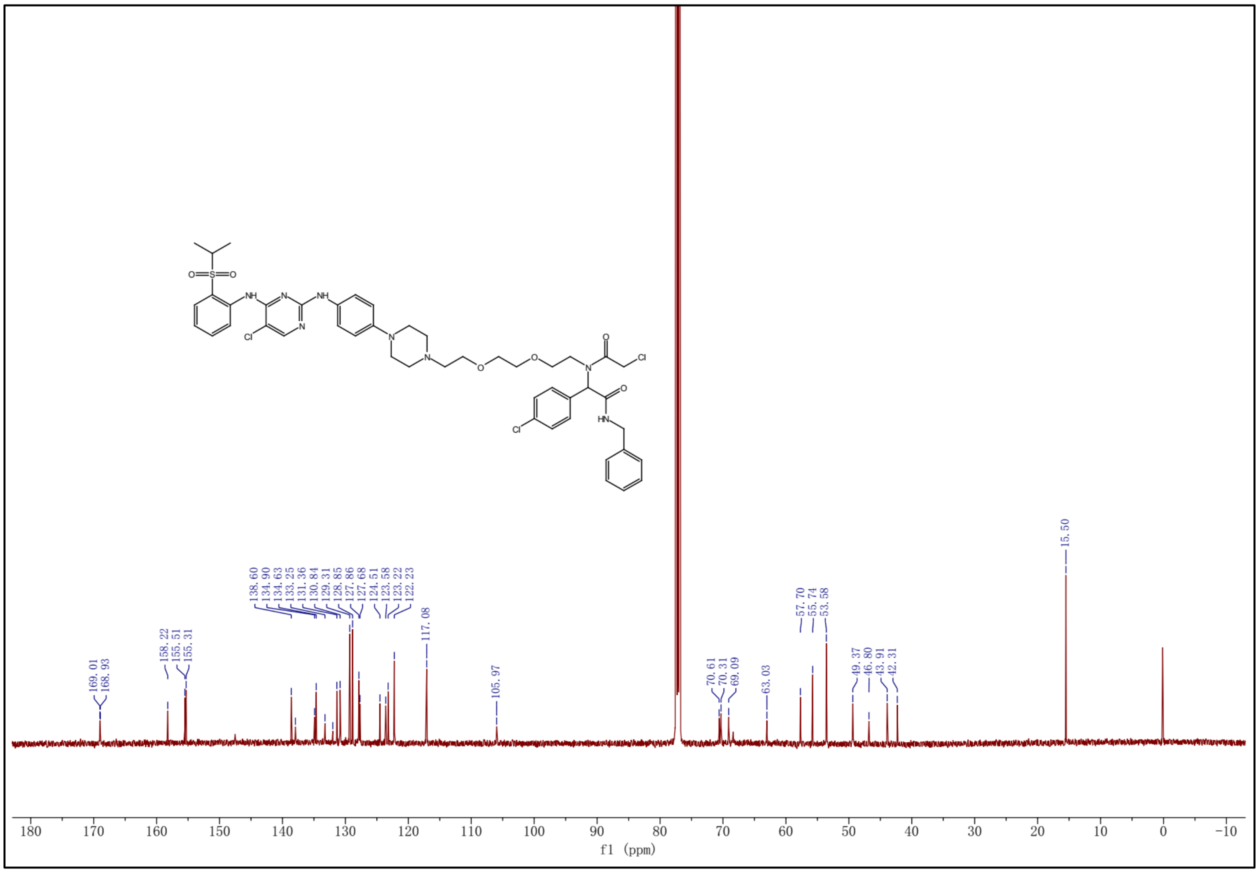

**Supplementary Figures**

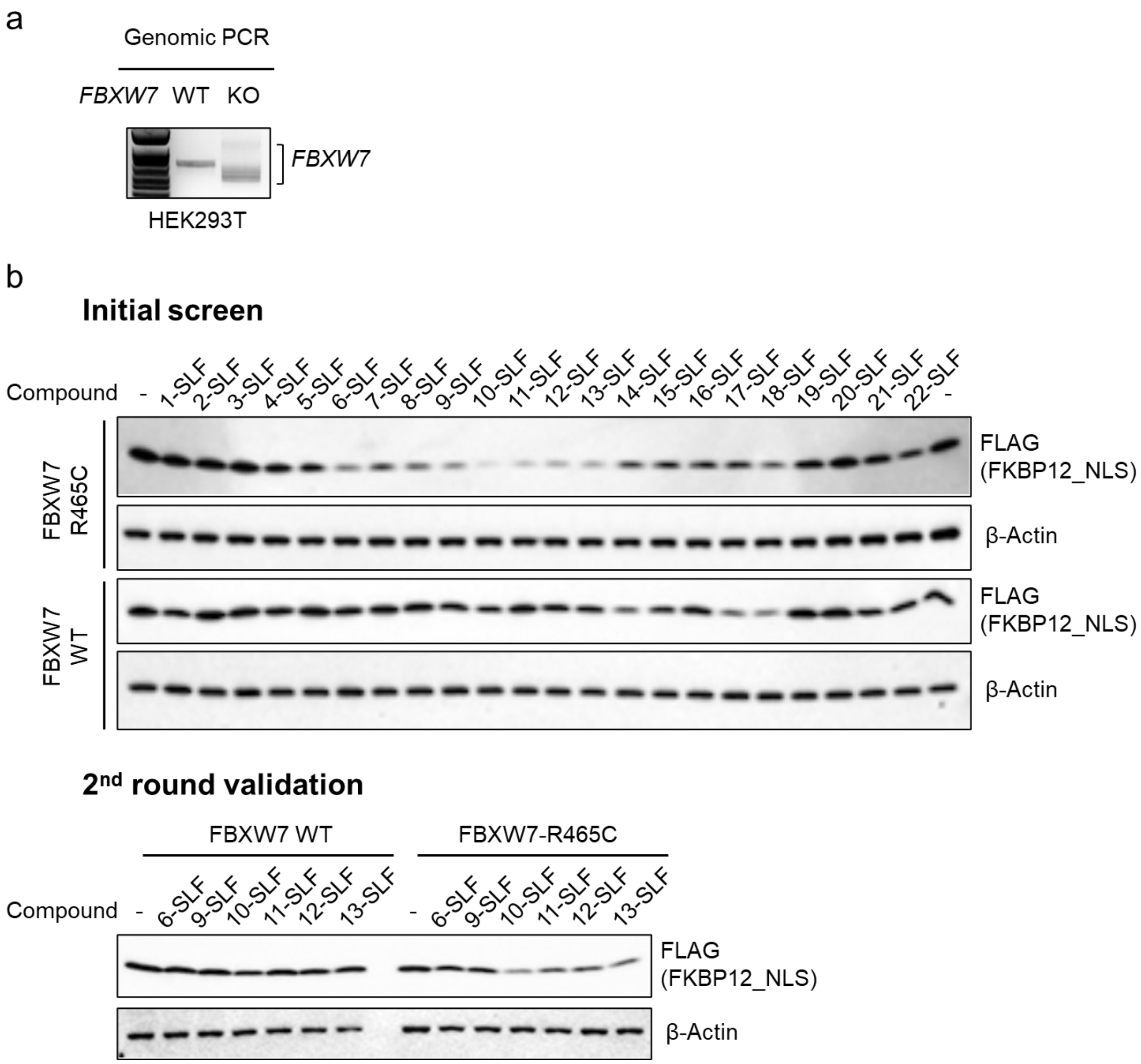


**Figure S1. Identification of heterobifunctional compounds that degrade FKBP12_NLS in an FBXW7-R465C-dependent manner**. **a**. Genomic PCR confirms *FKBW7* KO in HEK293T cells. **b**. Western blot analysis of a focused FKBP12-directed heterobifunctional compound library to identify compounds that induce FKBP12_NLS degradation in an FBXW7-R465C-dependent manner.

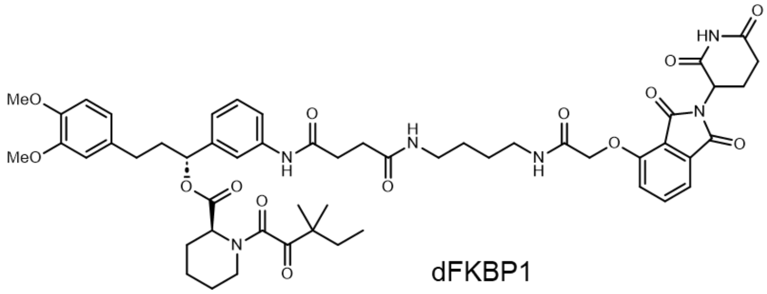


dFKBP1

**Figure S2.** Structure of dFKBP1.

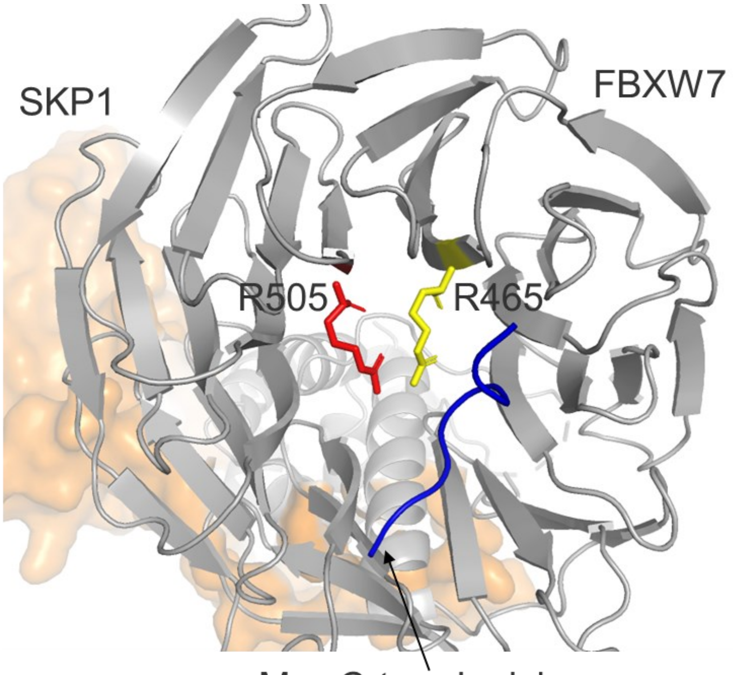


**Figure S3.** The crystal structure indicates that R505 is positioned near R465.

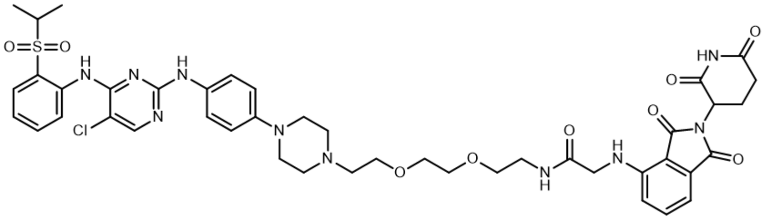


D1

**Figure S4.** Structure of D1.
